# Mechanistic Interpretability of Protein Language Models Reveals Encoded Structural and Functional Properties of Intrinsically Disordered Proteins

**DOI:** 10.64898/2026.09.10.750691

**Authors:** Lauren Naworski, Lydia L. Good, Rob Scrutton, Tuomas P. J. Knowles

## Abstract

Protein language models (PLMs) such as ESM-2, trained with a masked objective that leverages evolutionary constraints, encode protein sequences as embeddings for downstream tasks. Interpretability studies of ESM-2 have focused on folded proteins, leaving its behavior on intrinsically disordered proteins (IDPs) under-studied, despite IDPs constituting a substantial fraction of the human proteome and being implicated in numerous diseases. Because IDPs experience different evolutionary constraints, we hypothesized that PLMs would behave differently on disordered versus folded regions. Here we show that ESM-2 exhibits reduced attention on disordered regions, yet still encodes meaningful biological signals. The model assigns heightened attention to disease-relevant residues even at high levels of disorder. Both the radius of gyration and individual dynamic contact maps, key characteristics of IDPs, can be obtained from the model logits and embeddings. These findings suggest PLMs capture valuable information about IDP biology despite their bias toward structured residues.

## Introduction

Protein language models (PLMs) are usually transformer-based models (*1* ) trained on protein sequences with a masked learning objective, utilized to embed protein sequences for use in downstream tasks such as function, structure or interaction prediction (*2* ). By learning the rules of protein sequence grammar, PLMs not only learn about sequence patterns, but also the corresponding structural and functional traits of proteins inherently tied to their sequences, which are shaped by evolution (*2* ).

One such widely utilized transformer model is ESM-2, which is trained on an unsupervised masked learning objective across millions of protein sequences, capturing residue patterns that emerge through evolution to achieve its training objective (*3* ). While studies of ESM-2’s behavior and learned biological and physical information have been reported, these studies have focused on folded proteins. By nature of their structural plasticity, intrinsically disordered proteins (IDPs) exhibit different biological and physical behaviors (*4* ) as well as distinct sequence patterns (*5* ) compared to their folded counterparts. Given the flexibility of IDPs, there is less evolutionary constraint on mutations to maintain 3-D structure compared to folded proteins (*4* , *6* ). Because IDPs exhibit less evolutionary conservation than their folded counterparts, we hypothesize that PLMs will exhibit different behavior for disordered regions, specifically that their masked learning objective may bias them towards easier-to-predict, more highly-conserved folded domains.

IDPs and structured proteins including significant levels of disorder regions (IDRs) comprise up to 60% of the human proteome (*7* ). These proteins also play pivotal roles in numerous pathologies such as cancers and neurodegenerative diseases when their functions are disrupted by mechanisms such as misfolding or aggregation (*8* ). Given their abundance and relevance in human pathologies, these proteins represent a vast area for scientific advancement, and PLMs might be learning information necessary to predict features, structure, or functions of these proteins for use in downstream tasks such as targeting disease pathways. Before looking to advance PLMs’ application to IDP behavior, we must first establish an understanding of what IDP-specific information PLMs contain.

Early computational studies of IDPs utilized their distinctive sequence features to distinguish disordered from folded regions in protein sequences. The IUPred2a method relies upon energy estimation based predictions (*9* ), while the *D*^2^*P* ^2^ method and Metapredict combine predictions from many existing disorder predictors (*10* , *11* ). These computational methods have helped to shed light on the prevalence of disorder. In addition to predicting disorder, methods utilizing PLM embeddings have been used to predict certain properties pertaining to IDPs, such as IDP binding propensity (*12* ). While these tools predict disorder and properties of IDPs such as binding propensity, less is known about how modern PLMs understand, attend to, and represent disordered regions.

While previous work elucidating the behavior of PLMs for specifically disordered regions has not been reported, prior studies have explored ESM-2 embedding representation behavior for sequences at large, suggesting ESM-2 is capable of learning beyond sequence pattern. For example, analysis of an ESM-2-derived categorical Jacobian showed that ESM-2 stores motifs of pairwise inter-protein contacts (*13* ). This study focused on protein sequences with experimental structures missing at most 50 residues, thus eliminating disordered proteins from its dataset. Other work has revealed that latent features of PLMs capture biophysical features such as catalytic sites, zinc fingers, mitochondrial targeting sequences, phosphorylated residues, disulfide bonds and disordered regions (*14* ). Although this work identified features associated with disorder, it does not uncover how the model learns about disorder, only that the model can identify disorder. Interpretability studies have also directly examined attention of ESM-2, finding high attention sites linked to protein family classification (*15* ) and revealing patterns such as attention targeting binding sites and certain heads targeting specific amino acids (*16* ). Attention-based studies have hitherto not explored attention specifically for disordered proteins, which we anticipate to show differing attention patterns on account of the differences in the evolutionary constraint between disordered and folded proteins.

Given the lack of studies on PLM representation of disordered regions, combined with our hypothesis that PLM training objectives bias models against disorder, we seek to assess what information PLMs learn about IDPs through a study of PLM performance, logit, attention, and embedding analysis explicitly on disordered protein sequences. We first establish that differences in model behavior between folded and disordered regions exist through comparative analysis of differences in embedding representations, model performance, and attention. We next explore these differences by a comparative attention analysis, elucidating how the model attends differently in disordered versus ordered sequence contexts. We then extend our study to explore the ESM-2 signal for residues important to IDP structural ensembles and function, such as those involved in protein contacts and pathologically-relevant residues. With evidence of the model encoding structural information for IDPs, we show a multi-layer perceptron trained on PLM embeddings of IDP sequences achieves good performance in predicting IDP radius of gyration (*R_g_*). Our study sheds light on the previously unreported area of PLM interpretability for IDPs, finding that PLM training biases language models against disorder but retains valuable biological information pertaining to IDPs. This understanding will help inform future modeling strategies to better optimize PLMs for down-stream tasks related to disordered proteins.

## Results

We theorize that the differences between folded and disordered proteins, such as differences in amino acid composition (*5* ), evolutionary constraints required to maintain function (*6* , *17* ), and differences in the nature of intra-and inter-protein contacts (*18* ), may yield meaningful differences in PLM representations of the two groups, and correspondingly differences in the information stored in the model to make such representations.

To determine whether ESM-2 learns distinct representations of intrinsically disordered versus folded proteins, we performed comparative analyses of two matched datasets: disordered proteins from DisProt (*19* ) and folded proteins from the Protein Data Bank (PDB) (*20* ) (see Methods). While we utilize a subset of the sequences from both DisProt and PDB, we will refer to our filtered, matched datasets utilized in this work as “PDB” and “DisProt” proteins in our analyses.

### Embedding representations of folded and disordered proteins are partially separable and highly classifiable

We first compared ESM-2 output embedding representations between PDB (n = 2,120) and DisProt (n = 2,120) proteins to see if the model learns distinct representations for folded versus disordered sequences.

We examined the separability of the standardized final layer mean-pooled ESM-2 embeddings for the PDB and DisProt proteins via UMAP (*21* ) dimension reduction (see Methods). Dimension reduction revealed that a large cluster of DisProt proteins forms a distinct group apart from PDB proteins (Figure 1A, left). The partial overlap is expected, as our sample proteins contain regions of varying disorder (Figure S1B). The spatial distribution of sequences by IDR fraction (Figure 1A, right) largely aligns with the spatial distribution by dataset grouping and points to overlap on account of mixed degrees of disorder. The silhouette score of the original 1,280 dimensional embeddings (0.158) suggests weak separation on account of disorder.

**Fig. 1:**
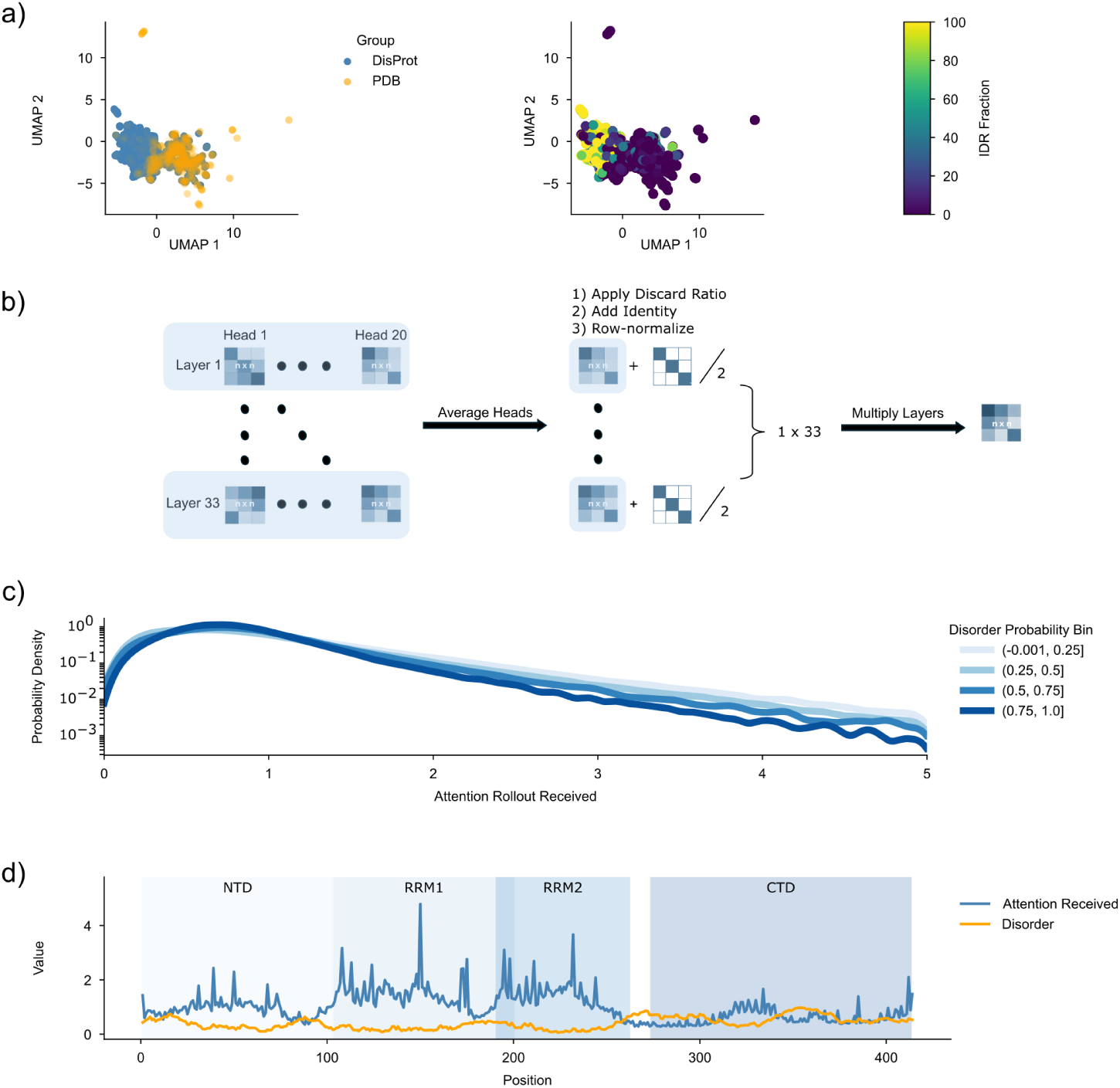
ESM-2 attention rollout shows a negative association to disorder. **A)** (Left) UMAP dimension reduction (*n_neighbors_* = 15, *min_dist_* = 0.1) of PDB (n = 2,120) and DisProt (n = 2,120) protein sequences’ scaled ESM-2 1,280 dimensional embeddings reveals partial overlap, with a subset of DisProt proteins forming a distinct cluster, suggesting differences in model embedding representations of folded and disordered proteins. (Right) The same UMAP analysis is shown where each sequence is colored by the fraction of sequence in a disordered region. This reveals the most-folded and most-disordered proteins cluster furthest from each other, with a gradient of disorder constituting the middle overlapping regions. **B)** Attention rollout (*22* ) involves averaging attention matrices across heads, and then successively multiplying the head-averaged matrices throughout the layers, resulting in one [Sequence Length, Sequence Length] attention rollout matrix. **C)** The kernel density estimates of attention rollout received by the residues of different IUPred2a (*9* ) disorder probability bins for residues in PDB and DisProt proteins reveals broadly similar distributions with the highest density at lower attention values, with the most structured bins having the widest distributions and largest tails of higher attention. **D)** Attention rollout received and IUPred2A (*9* ) disorder probability across each residue in TDP-43 exemplifies the inverse relationship between attention received and disorder. Attention received decreases notably in the heavily disordered C-terminal domain (CTD) (*23* ).

Considering UMAP cannot perfectly preserve the original structure of the embeddings, we complemented our UMAP analysis by assessing classification utilizing the 1,280 dimensional embeddings. A logistic classifier achieved 89.9% accuracy in classifying DisProt and PDB proteins (5-fold supervised cluster-stratified cross-validation, CD-HIT (*24* ) 40% similarity threshold). This suggests that the embedding representations of folded and disordered proteins are highly distinguishable.

Finding partial spatial separability and highly-distinguishable classification reveals evidence of differing model embedding representations between folded and disordered proteins.

### ESM-2 perplexity increases with disorder

With evidence of differing model representations for DisProt and PDB proteins, we would similarly anticipate differences to arise in model performance. Given that folded proteins exist in stable constructs with static contacts, whereas IDPs exist in flexible conformations with transient, low-affinity contacts (*25* ) more prone to mutation (*6* , *17* ), we anticipate the masked learning objective to be more difficult for the model in disordered regions, where there is more possible sequence variability and less evolutionary constraint. We evaluated the association of model performance and disorder by linear regression analysis (see Methods).

We calculated the ESM-2 perplexity of each DisProt and PDB protein (n = 4,240) utilizing a masking rate of 15%, according to the masking rate of the ESM-2 training (*3* ). We next regressed perplexity onto the mean disorder probability (as calculated by IUPred2a (*9* )) of the masked residues. The regression yielded a statistically significant positive coefficient (*β* = 6.20, *CI_lower_* = 5.35, *CI_upper_* = 7.05, *p* = 3.7 *∗* 10*^−^*^46^, *r*^2^ = 0.0496, *T_stat_* = 14.26, standardized *β^∗^* = 0.223) on masked residue disorder. These results imply a statistically significant weakly positive association of model perplexity to disorder, suggesting the model performance worsens on disordered regions.

This finding reinforces our hypothesis that PLMs are biased towards more stable, easier to predict folded regions on account of their training objective. This also further necessitates a deep understanding of what information is being learned about IDPs by PLMs to evaluate their efficacy in spite of the observed reduced performance. To better understand how the model processes disordered regions differently, we next examined ESM-2’s attention mechanisms directly.

### ESM-2 attends less to disordered regions of proteins

Given that ESM-2 shows reduced performance on disordered regions, we hypothesized that this performance deficit might manifest in altered attention patterns. Specifically, if the model struggles to predict disordered residues due to their higher sequence variability evolutionarily (*6* , *17* ), attention weights might systematically differ between ordered and disordered regions.

Attention rollout approximates the cumulative propagation of attention by combining attention heads across each layer and then successively multiplying the combined attention matrices from the first layer to the last (*22* ) (see Methods). This results in one row-normalized [Sequence Length, Sequence Length] attention rollout matrix.

Each row sums to 1 and row values represent the attention given by the residue at that row index across tokens of the sequence. Summing columns thus represents the total attention received by each residue at that column index after information propagation. We refer to this attention as *attention rollout received* to distinguish from the raw attention received at each head and layer. This method allows us to compare attention patterns in lower dimensions, where each protein sequence has 1 attention received value per residue. This calculation is shown illustratively in Figure 1B for rollout of ESM-2’s 20 heads and 33 layers.

To test our hypothesis that attention weights might differ between folded and disordered regions on account of sequence variability differences, we assessed if there exists any relationship between attention received and disorder. For PDB and DisProt proteins, the probability densities of residue-level attention rollout received grouped by differing disorder probability bins revealed highly-disordered residues to see lower probability densities at high attention (Figure 1C). All distributions see the highest density at low-attention values. The most disordered bins see the highest peaks, suggesting the tightest distributions with residues in these bins being more closely centered around low-attention. The most structured bins see the widest distributions, and correspondingly the highest density at the high-attention tail.

This pattern is evident in TDP-43, a well-characterized IDP associated with amyotrophic lateral sclerosis, frontotemporal dementia, and Alzheimer’s disease (*23* ). The protein’s C-terminal prion-like domain (CTD)(residues 274-413), which is highly disordered, receives markedly less attention than the structured N-terminal (NTD) and RNA recognition motifs (RRM1, RRM2) (Figure 1D). See Supporting Information for further examples (Figure S2).

We intuit the under-attention of disordered regions again based on the training objective of the model being biased for stable, folded regions. Folded domains exist in stable, high affinity contacts (*18* ) with highly conserved motifs, while disordered proteins experience more mutability and correspondingly higher sequence variability (*6* , *17* ), offering less predictive value. To achieve its masked learning objective, the model learns in training to attend more highly to the more predictable folded domains. While disordered regions might be less-attended to by PLMs, these models might still be learning valuable information with biological relevance for IDPs. We therefore examined whether the attention patterns of folded and disordered proteins reveal biologically interpretable differences in how ESM-2 processes specific amino acids in disordered contexts.

### ESM-2 mildly distinguishes amino acid attention allocation in disordered contexts

We have shown that ESM-2 under-attends and under-performs in disordered regions relative to folded regions, but this finding does not exclude the possibility of biologically-relevant information being encoded for IDPs. Expanding from our findings revealing that ESM-2 handles folded and disordered regions differently, we next explored *how* ESM-2 handles folded and disordered proteins differently. We comparatively examined the attention allocation to each amino acid type across PDB and DisProt proteins to evaluate the biological intuition in disordered contexts. We examined attention patterns through attention rollout (*22* ) to capture the cumulative attention representation, as well as raw attention at each head and layer.

We averaged the attention rollout received by residues across amino acid identity for PDB and DisProt proteins (Figure 2A). Average allocation across the amino acids was relatively similar between the two groups. Given the model is trained upon sequences in the UniRef database (*26* ), consisting of a spectrum of disordered regions, we would not anticipate the model to show drastically different attention allocation between folded and disordered proteins on average. Rather, what we observe is more moderate distinguishing of allocation, especially for biologically-key residues, depending on the disordered or ordered contexts.

**Fig. 2:**
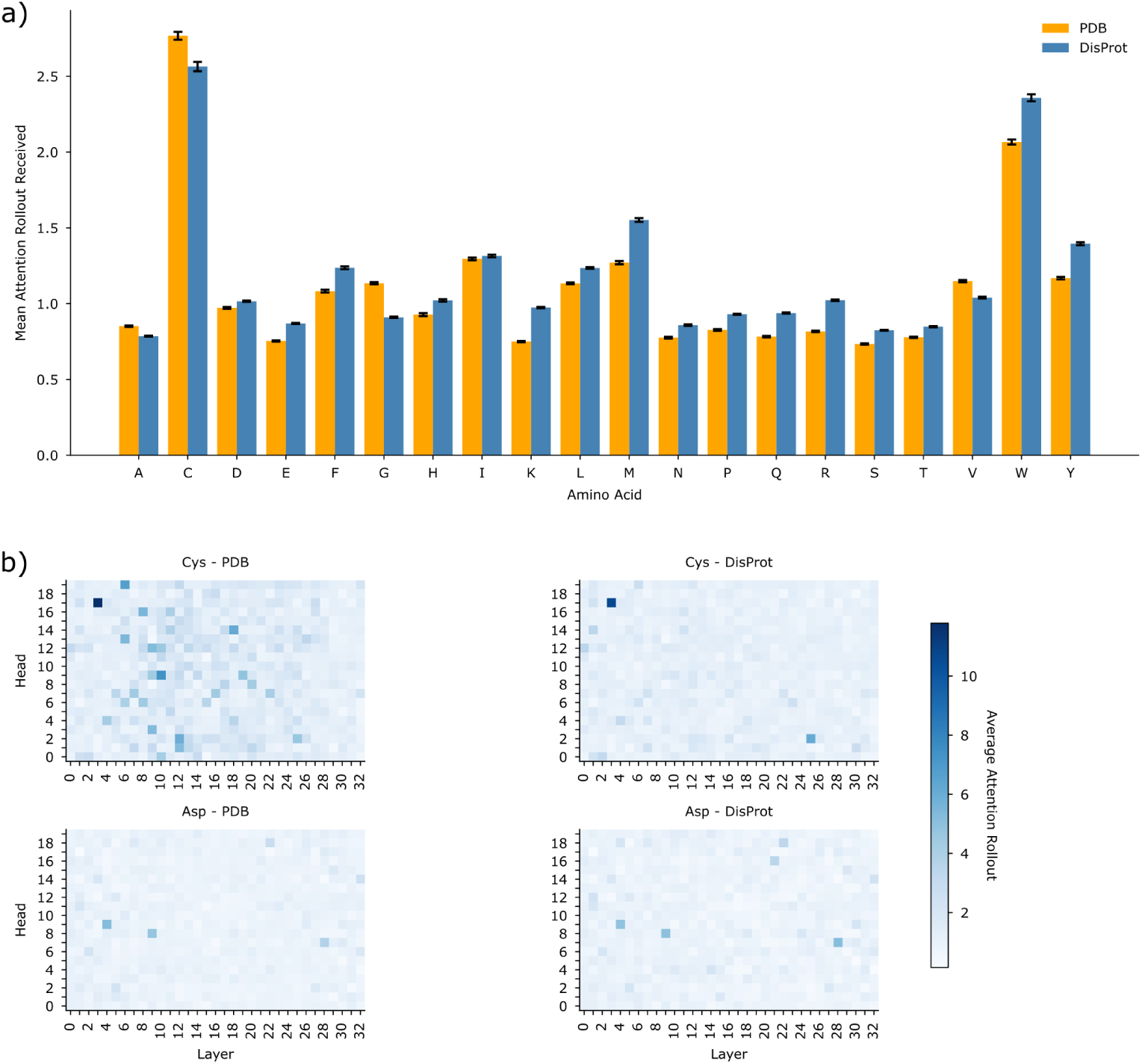
ESM-2 allocates similar amounts of attention on average to the same amino acids. **A)** The attention rollout received by each type of amino acid was averaged across all residues in either the DisProt (n = 2,120) or PDB (n = 2,120) subsets. Both PDB and DisProt proteins on average attended each amino acid with a similar level of attention, but amino acids such as Cys and Trp saw larger differences. Standard error of attention received across all residues in either PDB and Disprot are shown in black bars. Cohen’s *d* effect sizes are reported in Table 1. **B)** For each PDB and DisProt protein, attention received by each amino acid type was summed across all instances in the sequence at each head and layer, and divided by the number of occurrences of that amino acid type in the sequence. This occurrence-normalized sum was then averaged across all proteins in the PDB and separately in the DisProt sets. The average of this summed attention received normalized by occurrence is shown by color scale at each head and layer. Some amino acids such as Asp had highly similar attention patterns, but others such as Cys had large differences. Cohen’s *d* effect sizes and t-test p-values are reported in Table 2.

**Table 1:** Effect size (Cohen’s *d*) and independent Welch’s t-test p-values of DisProt and PDB attention rollout received distributions across amino acids (Cohen’s *d* subtracting Disprot - PDB).

| Amino Acid | Cohen's *d* | *p*-value | *n*<sub>DisProt</sub> | *n*<sub>PDB</sub> |
|---|---|---|---|---|
| G | -0.249 | 6.589e-162 | 24590 | 23882 |
| V | -0.117 | 1.331e-32 | 15207 | 21469 |
| A | -0.102 | 5.338e-31 | 24819 | 25740 |
| C | -0.102 | 4.632e-07 | 2779 | 6747 |
| I | 0.019 | 1.026e-01 | 10037 | 16793 |
| D | 0.063 | 2.491e-09 | 19051 | 17354 |
| L | 0.116 | 7.536e-39 | 20971 | 28209 |
| H | 0.119 | 6.607e-14 | 6199 | 8500 |
| T | 0.128 | 2.869e-35 | 17930 | 19334 |
| N | 0.135 | 1.514e-28 | 13384 | 13510 |
| S | 0.166 | 6.961e-71 | 31488 | 21465 |
| F | 0.169 | 5.047e-33 | 8035 | 11712 |
| P | 0.182 | 5.495e-58 | 24556 | 14153 |
| E | 0.217 | 5.295e-117 | 26540 | 21030 |
| W | 0.274 | 2.183e-24 | 2046 | 4289 |
| Y | 0.274 | 2.786e-72 | 6684 | 10908 |
| Q | 0.286 | 4.849e-126 | 16305 | 12845 |
| M | 0.304 | 5.540e-65 | 5790 | 7168 |
| R | 0.338 | 9.575e-205 | 16237 | 16905 |
| K | 0.383 | 0.000e+00 | 20458 | 19753 |

**Table 2:** Summary statistics of Cohen’s *d* (Disprot - PDB) values comparing summed attention received normalized by occurrence distributions at each head and layer for DisProt and PDB proteins. Summary statistics are calculated only with data when head/layer/amino acid/PDB or DisProt Grouping combinations have false-discovery rate adjusted Welch’s independent t-test p-values values *<* 0.05.

| Amino Acid | Count | Mean | Std | Min | 25% | 50% | 75% | Max |
|---|---|---|---|---|---|---|---|---|
| G | 611.0 | -0.314 | 0.386 | -1.256 | -0.567 | -0.360 | -0.146 | 1.112 |
| C | 595.0 | -0.288 | 0.266 | -0.846 | -0.453 | -0.351 | -0.206 | 0.901 |
| V | 609.0 | -0.217 | 0.417 | -1.278 | -0.464 | -0.263 | -0.072 | 1.362 |
| A | 552.0 | -0.211 | 0.378 | -1.204 | -0.445 | -0.256 | -0.066 | 1.118 |
| L | 561.0 | -0.137 | 0.489 | -1.502 | -0.457 | -0.162 | 0.196 | 1.357 |
| I | 560.0 | -0.103 | 0.405 | -1.141 | -0.347 | -0.135 | 0.165 | 1.315 |
| W | 477.0 | -0.052 | 0.317 | -0.915 | -0.280 | -0.122 | 0.176 | 1.005 |
| D | 569.0 | -0.031 | 0.465 | -1.449 | -0.313 | -0.096 | 0.266 | 1.477 |
| H | 471.0 | -0.013 | 0.333 | -0.855 | -0.231 | -0.095 | 0.190 | 1.143 |
| P | 588.0 | -0.007 | 0.520 | -1.183 | -0.362 | -0.100 | 0.343 | 1.584 |
| F | 539.0 | -0.002 | 0.410 | -1.103 | -0.263 | 0.071 | 0.269 | 1.339 |
| S | 574.0 | 0.023 | 0.531 | -1.478 | -0.343 | 0.086 | 0.392 | 1.679 |
| T | 576.0 | 0.034 | 0.405 | -1.328 | -0.246 | 0.089 | 0.325 | 1.139 |
| N | 555.0 | 0.036 | 0.375 | -1.050 | -0.221 | 0.081 | 0.269 | 1.291 |
| Y | 541.0 | 0.076 | 0.382 | -0.901 | -0.177 | 0.140 | 0.306 | 1.380 |
| E | 589.0 | 0.116 | 0.501 | -1.472 | -0.225 | 0.159 | 0.448 | 1.317 |
| M | 537.0 | 0.117 | 0.282 | -0.882 | -0.083 | 0.157 | 0.259 | 1.096 |
| Q | 585.0 | 0.142 | 0.419 | -1.271 | -0.148 | 0.180 | 0.391 | 1.385 |
| R | 589.0 | 0.172 | 0.458 | -1.330 | -0.143 | 0.231 | 0.464 | 1.389 |
| K | 597.0 | 0.251 | 0.488 | -1.320 | -0.118 | 0.314 | 0.587 | 1.461 |

Cys residues received the highest average attention in both groups. While Cys might not be engaged in disulfide bridges in disordered contexts, the model still learns in training that these residues are highly relevant and thus attends highly to them in both PDB and DisProt. However, DisProt proteins allocated less attention on average to Cys relative to PDB proteins. We reason that Cys is less likely to play its traditional structural role in disordered regions, prompting the model to allocate less to these residues in disordered context. To quantify the effect-size of differences in the distributions of attention to each amino acid, we report the Cohen’s *d* in Table 1, subtracting PDB from DisProt. While all of the amino acids saw statistically-significant differences, the effect-sizes were mostly small. Of the larger positive Cohen’s *d* values, suggesting DisProt proteins had a higher mean, were known disorder-promoting residues (Lys, Arg, Met) (*27* ) or residues involved in mediating IDP interactions (Trp) (*28* ).

Instead of analyzing attention through the cumulative rollout method, we next examine differences in attention patterns of each amino acid at every head and layer of the model to evaluate differences in how the model is organizing information. We calculated the total attention received by each amino acid per protein (summing attention across residues in the sequence by amino acid type) at each head and layer, normalized by amino acid occurrence in that protein sequence. We then averaged this summed attention per amino acid across the DisProt and PDB proteins at each head and layer. While some attention maps showed remarkably similar trends between the DisProt and PDB groups, such as in the case of Asp, other amino acids like Cys had larger differences (Figure 2B).

The contrasting attention patterns between PDB and DisProt proteins for Cys reveal context-dependent processing by the model. In DisProt proteins, Cys is primarily processed through few specialized attention heads, consistent with documented head specialization in transformers where certain heads focus on specific tokens or specific amino acids (*16* , *29* ). Analogous to induction heads, which specialize in in-context learning (ICL) and whose ablation significantly degrades ICL performance (*29* ), these specialized cysteine-focused heads may handle a singular, well-defined aspect of disordered Cys residues. In PDB proteins, however, the model allocates attention to Cys across multiple heads and layers simultaneously. This distributed attention pattern suggests the model processes structural Cys through multiple representational sub-spaces, potentially capturing their involvement in multiple concurrent functions such as disulfide bond formation, structural contacts, and folding constraints. The shift from specialized head attention in disordered contexts to distributed multi-head attention in folded contexts indicates the model has implicitly learned that Cys’ functional complexity varies with structural context.

To quantify the effect-sizes of differences in raw attention distributions of the amino acids between DisProt and PDB proteins, we calculated Cohen’s *d* at each head and layer for each amino acid by comparing the distributions across proteins of the previously described occurrence-normalized summed attention received per amino acid between PDB and DisProt (subtracting DisProt - PDB). 11,275 out of 13,200 Cohen’s *d* values were significant. We report the summary statistics of the Cohen’s *d* values by amino acid across heads and layers and proteins, only including significant Cohen’s *d* (Table 2).

The majority of amino acids saw negligible average effect sizes *< |*0.1| (Trp, Asp, His, Pro, Phe, Ser, Thr, Asn, Tyr). Gly (mean Cohen’s *d* = -0.314) and Cys (mean Cohen’s *d* = -0.288) saw larger negative effect-sizes, showing PDB proteins to be attending more to these residues on average across heads and layers. Lys (mean Cohen’s *d* = 0.251), Arg (mean Cohen’s *d* = 0.172), and Gln (mean Cohen’s *d* = 0.142) conversely attended more highly on average to DisProt proteins across heads and layers. These differences in raw attention by effect-size largely align with those reported from the aforementioned rollout method. The residues with larger effect-sizes additionally tended to be disorder-promoting residues (Lys, Arg, Met) (*27* ) while highly important structural residues such as Cys (*30* ) saw larger negative effect-sizes. These results underscore the presence of model differentiation in allocation of attention dependent upon disordered or folded contexts, and additionally suggest material biological information being learned about both disordered and folded contexts.

Through examining attention allocation to the different amino acids at both the cumulative attention rollout level, as well as the raw attention at each head and layer, we have shown that while PLMs show similar average attention allocations, there exist biologically-meaningful differences in cumulative average attention to amino acids and model processing, exemplified by diffuse versus specialized attention, dependent on disordered or ordered contexts. While ESM-2 shows degraded performance and less attention to disordered residues, these attention patterns reveal real biological signal being learned for IDPs.

### ESM-2 pays more attention to pathologically-relevant residues even at high degrees of disorder

With evidence of the model learning distinct biology for IDPs through distinguishing of attention allocation, we next explored if the model also learns information pertaining to the structure and functions of IDPs arising from their amino acid constituents. Given the training objective of ESM-2 relies upon the model learning the sequence grammar emergent through evolution (*3* ), the model learns the patterns of highly-conserved residues. These residues become highly conserved on account of their importance to protein folding (structure) or function (*31* ). We hypothesize that PLMs may learn information related to functionally and structurally critical residues of IDPs, which we evaluated via attention and logit analyses.

To see if the model encodes information pertaining to protein function, we examined ESM-2’s sensitivity to pathogenicity by comparing the attention received by disease-relevant residues to other residues. We utilized 3,356 unique protein sequences with documented pathogenic variants reported through ClinVar database (*32* ) (see Methods). For each protein, residues with known single-nucleotide pathological variants, tagged as “Pathogenic” or “Likely Pathogenic” by ClinVar database (*32* ), were labeled as “Disease-Relevant” and all other residues were labeled as “Non-Disease-Relevant.” Attention rollout was peformed on the wild-type sequences of each protein, resulting in an attention rollout received value for each residue. Comparing disease-relevant residues to non-disease-relevant residues, disease-relevant residues tended to fall in a higher attention percentile, where percentiles were calculated within each protein (Non-Disease-Relevant Median (IQR): 0.50 (0.25–0.75); Disease-Relevant Median (IQR): 0.68 (0.43–0.87); Figure 3A).

**Fig. 3:**
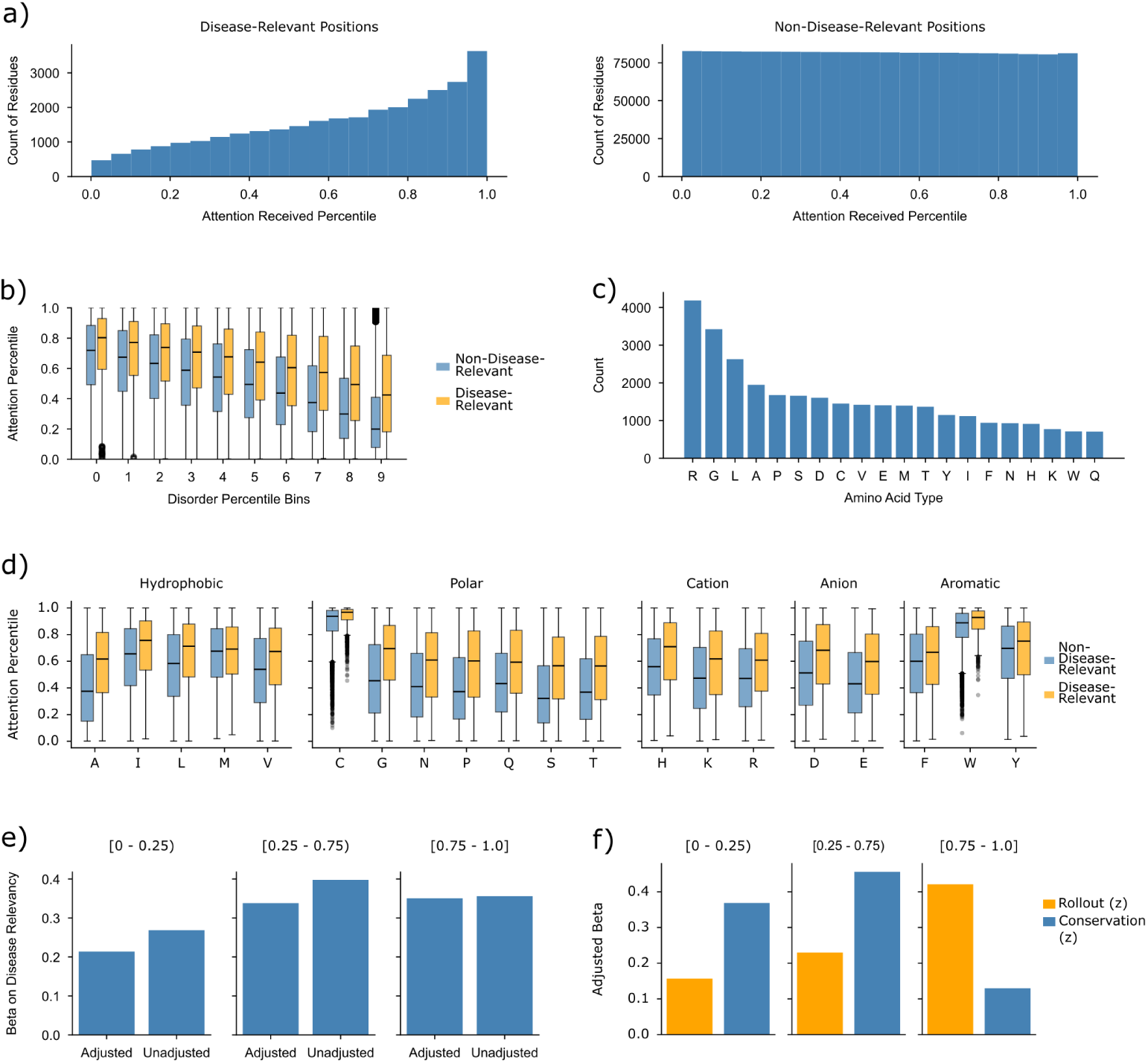
ESM-2 distinguishes residue pathogenicity through heightened attention to disease-relevant residues, even at high levels of disorder. **A)** Disease-relevant residues (Pathogenic or Likely Pathogenic in ClinVar (*32* )) from 3,356 proteins have higher within-protein attention rollout percentiles than non-disease-relevant residues (median [IQR]: 0.68 [0.43–0.87] vs. 0.50 [0.25–0.75]). **B)** Disease-relevant residues maintain higher attention percentiles across increasing disorder percentile bins. **C)** Disease-relevant residues are enriched in specific amino acids, with R, G, and L showing the highest counts. **D)** Attention percentile distributions are shown by amino acid for disease-relevant and non-disease-relevant residues. **E)** Attention rollout was regressed on disease relevance and DescribeProt conservation scores (*33* ) for residues of 3,015 ClinVar proteins, stratified by disorder probability bin. The unadjusted effect of disease relevance (*β >* 0.2) remains consistent across bins, while adjustment for conservation results in progressively less attenuation with increasing disorder. **F)** Logistic regression of disease relevance on standardized attention rollout and conservation shows that the adjusted coefficient for attention increases with disorder probability.

Given our prior finding of ESM-2 attending less to disordered regions, we explored if this phenomenon of heightened attention to disease-relevant residues was witnessed across varying degrees of disorder. When examining the attention percentile of both groups of residues by increasing levels of disorder (Figure 3B), attention decreases as disorder increases among both groups, as expected given our aforementioned findings. However, even as disorder increases, the median attention percentile of disease-relevant residues is still higher than that of the non-disease relevant residues, suggesting that the model is still distinguishing between pathogenicity to some degree even in highly disordered contexts.

The disease-relevant residues are not evenly distributed across amino acid identities, with certain amino acids like Arg, Gly, Leu being more highly populated amongst the disease-relevant residues (Figure 3C). We previously revealed certain residues, such as Cys and Trp, to receive higher attention rollout values on average than other residues. To ensure that the disease-relevant residues were not seeing higher attention on account of being disproportionately represented by high attention amino acids, we examined the attention percentile distributions of both disease-relevant and non-disease-relevant residues by each amino acid (Figure 3D). For the highest average attention rollout amino acids (Cys, Trp) the median attention percentile is roughly equal to that of the non-disease-relevant residues, revealing some of the smaller gaps in disease-relevant to non-disease relevant attention percentile medians across amino acids. This suggests that the disease-relevance effect is not a simple recapitulation of of amino acid-specific attention patterns.

Given our hypothesis that high evolutionary conservation of disease-relevant residues should direct the model’s attention toward these regions, we sought to quantify how much of the attention signal attributable to pathogenicity is explained by overlap with highly conserved residues. We regressed attention rollout onto disease relevancy and evolutionary conservation (using pre-computed conservation scores from DescribeProt (*33* )) across residues in 3,015 proteins, stratifying the analysis by residue disorder.

The *β* on disease relevancy showed the greatest attenuation upon controlling for conservation in the most-folded residue bin (*≈* 20%) (Figure 3E). This indicates that roughly 80% of the attention signal is independent of conservation, implying the model captures pathogenicity-associated features beyond evolutionary constraint. Attenuation diminished progressively across bins of increasing disorder, falling below 5% in the high-disorder bin, consistent with disordered regions being less evolutionarily-conserved.

To directly compare the predictive power of attention and conservation for residue disease relevancy, we regressed disease relevancy onto standardized attention rollout and conservation scores, again stratified by disorder bin (Figure 3F). Among the most-folded residues, the adjusted *β* on attention was less than half that of conservation. However, the *β* on attention rollout increased across the mixed- and high-disorder bins, substantially exceeding that of conservation for the most disordered residues, suggesting attention is a better predictor of disease-relevancy for disordered residues. These results reveal the model can capture patterns of highly-conserved motifs such as those connected to pathologies beyond conservation even at high degrees of disorder.

### ESM-2 signal correlates with IDP contact maps

Our finding of the presence of heightened attention to disease-relevant residues across degrees of disorder reinforces that PLMs, while still attending less to and performing worse for disordered regions, can still learn valuable information about IDPs, especially when that information is related to highly conserved residues. Expanding on our assessment of ESM-2’s sensitivity to highly-conserved residues, we move from functionally-critical residues such as disease-relevant residues, to structurally-critical residues, such as those involved in intra-protein contacts.

It has been shown previously that ESM-2 stores information on co-evolving residues of folded proteins via analysis of a categorical Jacobian matrix (*13* ). (*13* )’s method systematically mutated each residue in the sequence to all 20 possible amino acids and measured how each mutation perturbed the output probabilities (logits) for amino acids at all positions in the protein sequence, resulting in a coupling matrix of residue-residue contact strengths. While IDPs do not exist in a stable folded teritiary structure like their folded counterparts, they still exhibit interconverting, transient intramolecular interactions (*4* ), and can exhibit a spectrum of foldability rather than being entirely disordered, suggesting co-evolving contacts of IDPs might be recoverable via this Jacobian method as well. We repeat this Jacobian analysis for the sequences of the human intrinsically disordered proteome (IDRome) (*34* ) to evaluate if ESM-2 similarly learns the co-evolution of interacting residues in IDPs.

We repeated the categorial Jacobian calculation according to (*13* ) for sequences of the IDRome less than 160 amino acids length for computational efficiency (n = 19,960 out of 28,058 IDRs), retrieving residue-level coupling matrices with a Jacobian contact strength calculated for each residue pair. While there does not exist ground-truths of IDP contact maps on account of them lacking a stable tertiary structure, we approximated by utilizing contact maps derived from coarse-grained molecular dynamics (MD) (*34* ), resulting in a fraction of frames spent in contact for each residue pair (see Methods).

The observed spearman correlation of Jacobian contact strength and MD fraction in contact across the sequences showed a positively shifted distribution (median r = 0.2848, IQR: [0.1360, 0.3792]; Figure 4A).

**Fig. 4:**
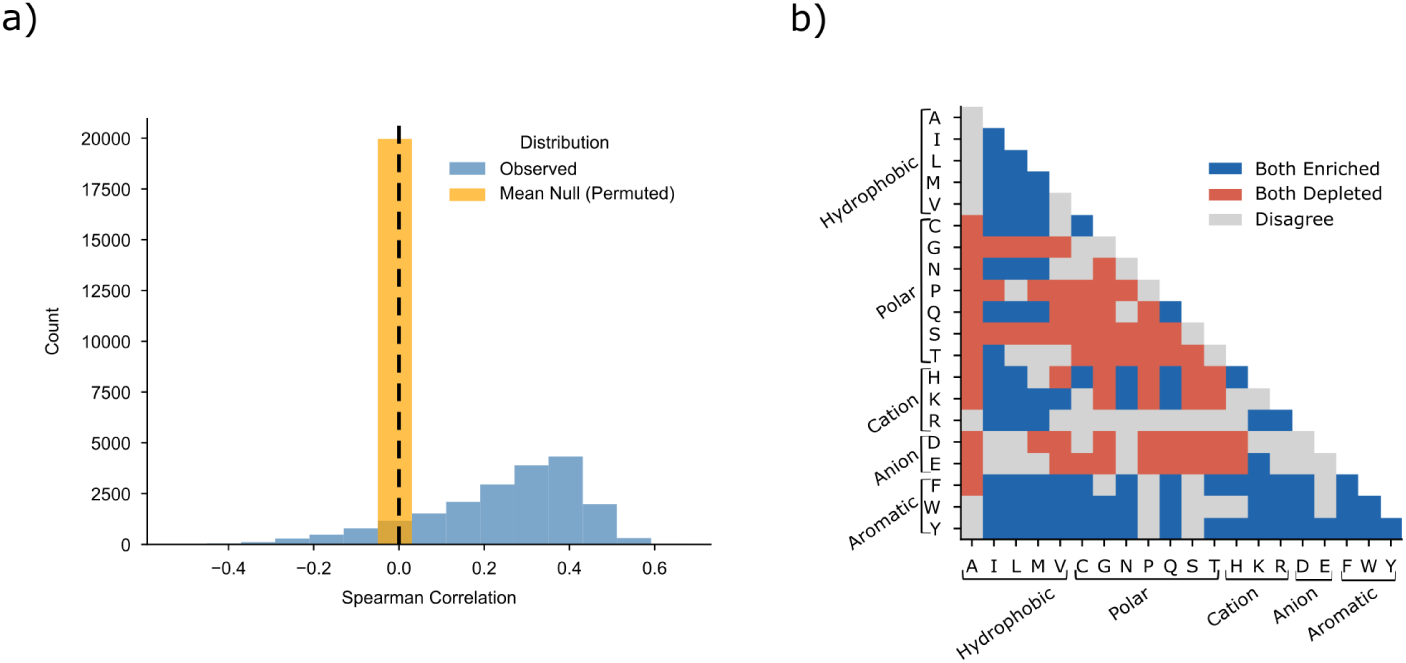
ESM-2 signal correlates with MD-derived contact maps of IDPs. **A)** ESM-2 derived categorical Jacobian matrices (*13* ) were reduced to residue-level coupling matrices of contact strengths for sequences of the human intrinsically disordered proteome less than 160 amino acids in length (*34* ) (n = 19,960). We calculated the spearman correlation of these Jacobian contact strengths with the corresponding MD-derived contact maps. Median spearman r = 0.2848, IQR: [0.1360, 0.3792]. Permutation tests of shuffling Jacobian contact strengths saw a mean null r = -0.00 ± 0.03, suggesting that the Spearman correlation of the Jacobian and MD contacts was far above null expectations. **B)** To assess the similarity of the two methods, we calculated the frequency of each amino acid pair in the top deciles of Jacobian and MD contact strength separately and compared this frequency to the baseline frequency across all residue pairs. The majority of amino acid pairs saw directionally the same enrichment or depletion across both methods.

To determine whether these observed correlations exceed chance expectations, we performed protein-specific permutation tests by randomly shuffling Jacobian contact strengths 100 times per protein while keeping MD-derived contact frequencies fixed.

The observed median Spearman correlation was far above null expectation (mean null r = -0.00 ± 0.03), corresponding to a median z-score of 11.33 across all proteins. Of 19,960 proteins analyzed, 83.1% (16,592 proteins) exhibited correlations above their null distributions (p *<* 0.05), suggesting that ESM-2 captures structural coupling beyond random chance.

While the effect size is modest (mean Δ*r* = 0.24, *r*^2^ *≈* 0.059), the signal is nevertheless meaningful given the intrinsic noise of MD-derived contact maps, which represent ensemble-averaged behavior simulated with a coarse-grained force field. This finding suggests that ESM-2 is storing a degree of signal relevant to co-evolving residues involved in intra-protein interactions of IDPs.

We compared the two methods by examining differences and similarities in which types of amino acid pairs comprised the top top decile of Jacobian contacts versus the top decile of MD contacts (Figure 4B). We calculated the frequency of amino acid pair in the top decile of Jacobian and MD contacts respectively, and we compared this to the baseline frequency across all residues. The majority of amino acid pairs saw the same directionality (enriched or depleted versus baseline) in both methods’ top deciles. This reinforces the similarity in representation between the ESM-2-derived and MD-derived contact maps. The raw magnitudes of enrichment and depletion for each pair and for each method are shown in Figure S3.

While we utilize MD-based estimations of IDP contact maps, the presence of non-random correlations between the evolutionarily-based and physical-based methods suggests that ESM-2 is storing signal of the co-evolving contacts of IDPs, capturing structural information even for these flexible proteins.

### ESM-2 Embeddings Predict *R_g_* of IDPs

With evidence of ESM-2 encoding signal to the structural properties of IDPs, we hypothesized that the model might be capable of efficaciously predicting properties pertaining to the conformations of IDPs, such as radius of gyration (*R_g_*).

We trained a multi-layer perceptron (MLP) upon the final-layer mean-pooled ESM-2 embeddings for sequences of the IDRome (*34* ) (n = 20,368) to predict *R_g_* after subtracting a length-dependent baseline to encourage the model to learn sequence composition signals beyond chain length (see Methods).

We compared our model to the reported MD-derived *R_g_* (*34* ) of the IDRome test set sequences (n = 5,444), wherein both *R_g_* values were normalized by the Flory scaling *R*_0_ *∗ N^ν^* to evaluate efficacy in our model predicting the expansion or compaction of sequences relative to the *R_g_* expected based upon sequence length alone (Figure 5A, left). The resulting correlation (Pearson R = 0.902, Spearman R = 0.880) suggests high model efficacy.

**Fig. 5:**
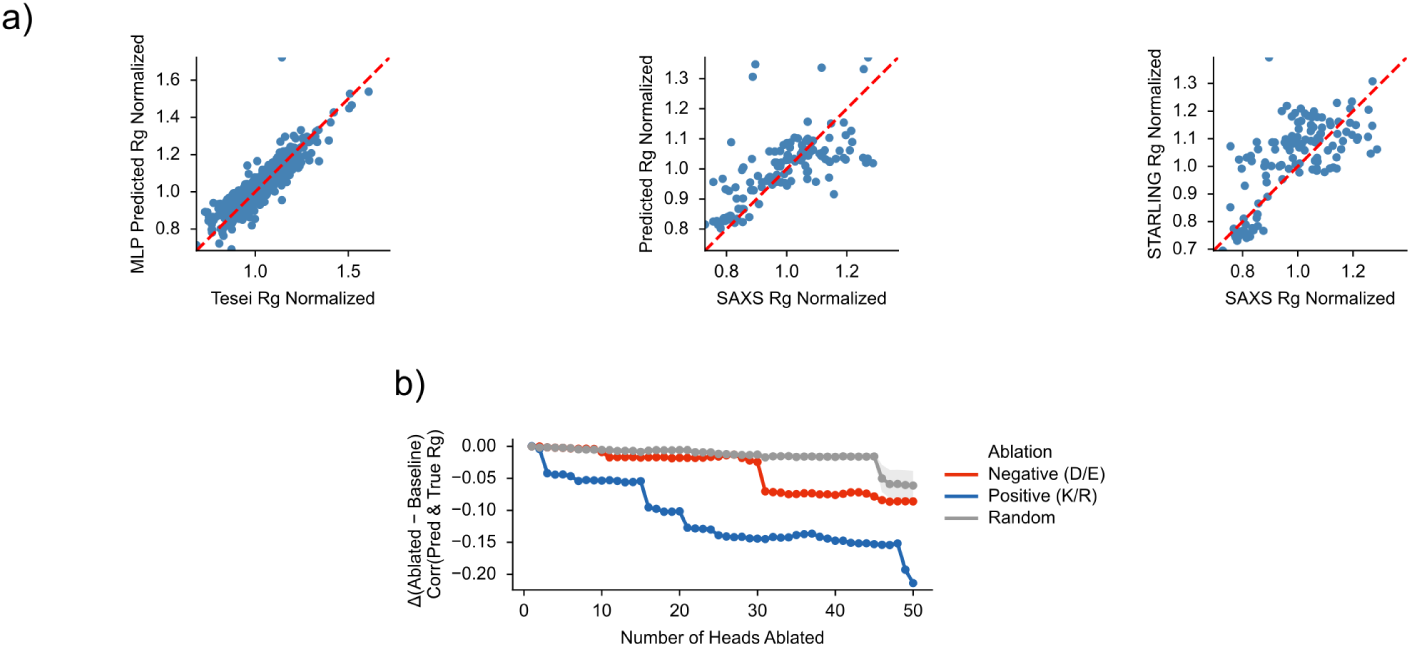
An MLP trained on ESM-2 embeddings accurately predicts radius of gyration (*R_g_*), and depends on charge-associated attention heads. A) Predicted versus ground-truth *R_g_*, each normalized by the empirical Flory scaling law (*R_g_/R*_0_*N^ν^*, with *R*_0_ and *ν* fit on the training set). Left: MLP predictions versus MD-derived *R_g_* for the IDRome test set (*34* ) (n = 5,444; Pearson *r* = 0.90). Middle: MLP predictions versus experimental SAXS-derived *R_g_* (*35* ) (n = 127; *r* = 0.68). Right: STARLING predictions (*35* ) versus the same SAXS-derived *R_g_*(n = 127; *r* = 0.69), showing performance comparable to the MLP. **B)** Change in prediction accuracy (Spearman *ρ* with MD-derived *R_g_*, IDRome test set) as attention heads are progressively zero-ablated. Heads were ranked by occurrence-normalized attention received by negatively charged (D/E; “Negative”) or positively charged (K/R; “Positive”) residues, averaged across DisProt and PDB proteins (n = 4,240). Targeted ablations are compared with ablating the same number of randomly selected heads (5 random permutations per number of heads).

We additionally tested the model efficacy compared to experimentally-validated *R_g_* values (n = 127) (*35* ), which saw a moderately reduced correlation (Pearson R = 0.677, Spearman R = 0.731, Figure 5A middle).

Our model shows similar performance to the existing STARLING model, which is a two-stage generative model that combines a variational autoencoder with a denoising-diffusion probabilistic model to create a latent diffusion model for predicting coarse-grained ensembles of IDPs (*35* ) (Pearson R = 0.691, Spearman R = 0.656, Figure 5A, right).

These results reveal high efficacy in utilizing ESM-2 for predicting *R_g_* of IDPs, achieving similar performance to the current leading models. The simple construct of our model combined with efficacious performance highlights the ability of out-of-the-box ESM-2 embeddings to be utilized for down-stream predictions pertaining to IDPs such as qualities of their conformations.

### ESM-2 Encodes Electrostatics for Charged Amino Acids

Considering our MLP was able to predict the expansion or compaction of IDPs relative to their expected Flory scaling with high efficacy, our model appeared to be learning conformational properties beyond just sequence length. We hypothesized that the ESM-2 embeddings might be encoding signals to electrostatic interactions, which could cause protein expansion or compaction due to charged repulsion or attraction, and thus allow our model to predict such expansion and compaction relative to Flory scaling.

To evaluate if ESM-2 encodes signal to electrostatic interactions, we ablated heads of the model we hypothesized to be highly involved in representing charged residues and compared the *R_g_* predictions of our *R_g_* MLP when given the ablated model embeddings versus the baseline model embeddings (see Methods).

To identify attention heads associated with electrostatic interactions, we examined the average attention received by each amino acid type at each head and layer of the model for all 4,240 PDB and DisProt proteins, and then averaged this across proteins. Attention allocation to the two negatively charged residues (Asp, Glu) was highly correlated with one another (r = 0.803), as were those for the positively charged residues (Lys, Arg) (r = 0.794) (Figure S5). Combinations of correlations between attention to oppositely charged residues were all low, suggesting that distinct subsets of heads preferentially attend to negative and positive charges respectively (r values: Asp, Lys (0.037); Asp, Arg(−0.096); Glu, Lys(0.276); Glu, Arg(0.085)). We hypothesized that the heads attending most strongly to acidic or basic residues are likely to encode electrostatic information.

We ordered heads by decreasing average attention to negatively charged and separately to positively charged amino acids and progressively ablated increasing numbers of heads up to 50 heads. We compared this targeted ablation to a control of random ablations wherein we randomly ablated the same number of heads replicated with 5 permuations of random ablations per number of heads ablated. For each ablation, we analyzed the difference in correlation of the Flory normalized MLP predicted *R_g_* and the ground-truth *R_g_* (MD-derived *R_g_* for IDRome sequences (*34* ) and SAXS-derived *R_g_* for experimental (*35* )) between the prediction based on the ablated model embeddings and the baseline embeddings.

We observed a larger decrease in correlation of predicted and ground-truth *R_g_* comparing baseline to ablated models for both the negative head and positive head ablations beyond that of the random control (Figure 5B). Both ablations also showed a statistically significant increase in absolute error (negative ablation MAE increase: +0.01 *±* 0.021; positive ablation MAE increase: +0.013 *±* 0.024) (Tables 3, 4). These results reveal that ablating attention heads which attend most to charged residues reduces the *R_g_* MLP’s ability to accurately predict *R_g_*expansion or compaction relative to Flory scaling, and suggest that these high-attention-to-charge-residue heads are partially responsible for representing features associated with electrostatic interactions. Ablation results of negative-associated heads and positive associated heads are reported respectively in Tables 3 and 4.

**Table 3:** Negative Charge (D/E) 50 Head Ablation Results. : Baseline vs. Ablated Model Performance (Correlations with Ground-Truth)

| **Dataset** | **Model** | **Spearman** | **Pearson** | **MAE (Å)** | **ΔError (FN)** | **% Worse** |
|---|---|---|---|---|---|---|
| IDRome | Baseline | 0.880 | 0.902 | 0.019 | +0.01 ± 0.021 | 67.5% |
|  | Ablated | 0.794 | 0.841 | 0.028 |  |  |
| SAXS | Baseline | 0.731 | 0.677 | 0.074 | +0.003 ± 0.017 | 57.5% |
|  | Ablated | 0.719 | 0.665 | 0.077 |  |  |

ΔError (FN) = per-protein mean error increase after ablation (Flory Normalised). One-sample *t*-tests vs. 0: IDRome *t* = 34.394, *p* < 0.0001; SAXS *t* = 2.310, *p* = 0.0225.

**Table 4:** Positive Charge (K/R) 50 Head Ablation Results. : Baseline vs. Ablated Model Performance (Correlations with Ground-Truth)

| **Dataset** | **Model** | **Spearman** | **Pearson** | **MAE (Å)** | **ΔError (FN)** | **% Worse** |
|---|---|---|---|---|---|---|
| IDRome | Baseline | 0.880 | 0.902 | 0.019 | +0.013 ± 0.024 | 70.6% |
|  | Ablated | 0.667 | 0.758 | 0.0313 |  |  |
| SAXS | Baseline | 0.731 | 0.677 | 0.074 | +0.004 ± 0.034 | 50.4% |
|  | Ablated | 0.703 | 0.630 | 0.0775 |  |  |

ΔError (FN) = per-protein mean error increase after ablation (Flory Normalised). One-sample *t*-tests vs. 0: IDRome *t* = 39.485, *p* < 0.0001; SAXS *t* = 1.282, *p* = 0.2021.

To test whether ablation removes information beyond sequence composition, we regressed the ESM-2 embedding on the corresponding charge fraction (frac*_DE_* or frac*_KR_*) and evaluated R*_g_*prediction from the residual embedding, which removes linear information on positively or negatively charged fraction content. For the negative (D/E-attending) heads, ablation reduced full-embedding performance substantially (Pearson *r*: 0.902 *→* 0.841), but this drop almost entirely disappeared after residualization (0.320 *→* 0.315; bootstrap gap = *−*0.005, 95% CI [*−*0.018, 0.007], *p* = 0.41), indicating that the effect of ablating these heads is explained by reduced sensitivity to D/E composition alone. For the positive (K/R-attending) heads, ablation produced a larger full-embedding drop (0.902 *→* 0.758) that was only partially attenuated after residualization (0.418 *→* 0.373; bootstrap gap = *−*0.045, 95% CI [*−*0.060*, −*0.031], *p <* 10*^−^*^4^), showing that roughly a third of the total effect reflects information beyond K/R content. These results indicate an asymmetry between the two head populations: ablating D/E-attending heads removes composition-explainable signal only, whereas ablating K/R-attending heads removes additional, composition-independent information relevant to R*_g_* prediction.

Together, these results suggest that while ESM-2’s representation of negative-charge electrostatics is largely reducible to raw negative or positive charge composition, its representation of positive-charge electrostatics captures additional, composition-independent information, consistent with the model encoding aspects of charge patterning beyond simple amino acid counts.

## Discussion

Through our systematic exploration of ESM-2 attention, logit, and embedding representations, we have presented a thorough investigation of PLM efficacy and learned information for IDPs. We have shown the model has degraded performance and lessened attention to disordered regions, reinforcing our hypothesis that ESM-2’s reliance upon evolutionary constraints on sequences might result in differing performance for disordered regions. While this is not unexpected, as we would anticipate folded proteins to have more highly conserved sequence patterns within which transformers can more easily predict masked residues, this finding necessitates a deeper understanding of what PLMs are learning about disordered proteins. We discovered meaningful patterns in attention both through cumulative attention rollout and direct head/layer attention which align with biological expectations. We saw evidence of ESM-2 holding signal for evolutionarily-conserved residues, such as pathologically-relevant and co-evolving residues. We expanded on this by showing this encoded information can be used to predict IDP conformation properties such as *R_g_*.

Our analysis sheds light on PLM behavior on IDPs. While previous interpretability studies of ESM-2 (*13* , *14* ) have revealed meaningful biological features in the model’s representations, these works either explicitly excluded disordered proteins from analysis (*13* ) or did not systematically examine whether the model treats IDPs differently from folded proteins (*14* ) . Our finding that ESM-2 exhibits degraded performance and reduced attention on disordered regions provides the first systematic characterization of this behavioral difference.

Our results both align with and extend prior findings. Consistent with (*13* ) ’s demonstration that ESM-2 captures inter-residue contact information, we find that ESM-2 embeddings correlate with MD-derived contact maps. However, we extend this finding to IDPs, showing that this signal persists even in highly disordered regions, and that ESM-2 embeddings can be used to predict *R_g_*. Similarly, while (*14* ) identified disorder-associated features through sparse autoencoders, our direct analysis of attention patterns reveals how the model differentially processes these regions through reduced but biologically meaningful attention allocation, and identifies specific heads associated with properties such as electrostatics. Together, these findings suggest that while PLMs are biased toward structured regions, they nonetheless encode valuable functional information about IDPs that extends beyond simple disorder prediction tools like IUPred2a (*9* ) or *D*^2^*P* ^2^ (*10* ).

We acknowledge limitations of our presented work. Our use of the attention rollout received metrics implies certain assumptions, and serves as a representation of attention information rather than a ground-truth. Our choice of averaging across heads, as opposed to other methods of combination such as minimum or maximum, transforms the raw attention data and potentially misrepresents attention weights when they are not normally distributed. The rollout method of matrix multiplication across layers additionally assumes that information is mixed linearly across layers (*22* ). While these are useful approximations for interpretability, allowing analysis in fewer dimensions, they transform the raw data and introduce implicit biases and assumptions.

Apart from simplifying calculations, we also acknowledge limitations across datasets utilized in this study. Our PDB and DisProt datasets used in comparative attention analysis do not represent fully disordered or fully folded structures, as they exist in a spectrum of disorder. In our analysis of assessing ESM-2 signal for coevolving residues of protein contact maps, we relied on comparison to contact maps approximated through molecular dynamics, which imports any biases in the molecular dynamics force fields in our comparison.

These results raise important questions about whether future PLM training strategies should explicitly account for disorder. Current masked language modeling may implicitly bias models toward structured regions due to their higher predictability. Alternative training objectives or data augmentation strategies that emphasize functional sites in IDPs could potentially improve model performance on disordered regions while maintaining strong performance on folded proteins. Methods to improve PLM handling of IDPs represents an important area for further work. Given the reliance on evolutionary constraints for masked learning objectives, finetuning methods utilizing alternative learning objectives, such as finetuning for downstream tasks pertaining to IDPs, could present an avenue for designing more effective IDP-focused PLMs.

While PLMs such as ESM-2 attend less to disordered regions and show worse performance on these regions, we have shown that their attention patterns support biological intuition and suggest material knowledge being held in the models about these proteins. We have additionally shown that the model stores a degree of signal for certain highly-conserved residues, even for less evolutionarily constrained IDPs, through our analysis of pathologically-relevant residues and residues of intraprotein contacts.

## Materials and Methods

ESM2 T33 650M UR50D was utilized for all analysis in this work.

### PDB and DisProt Dataset Creation

We filtered disordered proteins from DisProt (*19* ) for sequences between 50 and 1,024 amino acids in length (n = 2,120), referred to as the “DisProt” set of proteins in this work. A matched comparative dataset of folded proteins was gathered from Protein Data Bank (*20* ). PDB sequences containing X, O, and U were excluded. We then subsampled the remaining PDB sequences to match the number of samples in the DisProt set (n = 2,120), and to roughly match the DisProt dataset’s sequence length distribution (Figure S1A).

### Dimension Reduction

To assess similarities or differences in ESM-2 embedding representations comparing PDB (n = 2,120) and DisProt (n = 2,120) proteins, we reduced their standardized mean-pooled 1,280dimensional final layer embeddings utilizing UMAP (*n_neighbors_* = 15, *min_dist_* = 0.1) (*21* ).

### Linear Regression of Perplexity onto Disorder

ESM-2 perplexity was retrieved for each PDB and DisProt sequence (2,120 DisProt and 2,120 PDB proteins) by masking 15% of each sequence via Hugging Face (*36* ), utilizing the same masking rate as the original ESM-2 training (*3* ). The resulting perplexity was regressed onto the average IUPred2a (*9* ) disorder probability of the masked residues using heteroskedasticity-robust standard errors.

### Attention Rollout

We computed attention rollout according to (*22* ). We averaged attention weight matrices across heads and then successively multiplied the head-averaged matrices through each layer from 1 to 33, resulting in one sequence length by sequence length attention matrix per sequence. The matrix was row-normalized, such that each row contains the distribution of attention given by the residue at that row index, and the sum of columns represents the attention received by the residue at that column position from all residues in the sequences. Padding token such as < *cls* > and < *eos* > are not included in the rollout calculation as to only consider attention received from the other residues in the sequence. A discard ratio of 0.9 was applied prior to the matrix multiplication. Our study focuses on the column-summed attention rollout matrices, or the vector of attention received after attention rollout.

### Cohen’s *d* for Attention Heatmaps

For each protein in the PDB and DisProt sets, we measured ESM-2 attention received across each residue at each *head_j_* and *layer_i_*. We summed the total amount of attention received by each amino acid type across all residues in *protein_p_* at *head_j_* and *layer_i_*, and then normalized by the occurrence of each amino acid in *protein_p_*. We calculated Cohen’s *d* by subtracting the mean of PDB proteins’ summed attention received normalized by occurrence from the mean of DisProt proteins for each amino acid (DisProt - PDB), divided by the pooled standard deviation. Filtering for only head/layer/amino acid combinations which had false-discovery corrected Welch’s t-test p-values *<* 0.05, we evaluated the summary statistics of Cohen’s *d* across each head and layer for each amino acid (Table 2).

### Disease-relevant Dataset Creation

We retrieved human single-nucleotide variants from ClinVar (*32* ) (accessed February 3, 2026) and retained only those labeled as “Pathogenic”, “Likely pathogenic”, or “Pathogenic/Likely pathogenic”. Variants were mapped to protein sequences using the NCBI Gene gene2refseq table (*37* ) and RefSeq GRCh38 protein sequences (*38* ). We removed variants whose wild-type or mutant residue could not be mapped to a standard amino acid (e.g., nonsense variants annotated as “Ter”), and retained proteins longer than 50 and shorter than 1,024 amino acids, yielding 3,359 sequences. Residues of each sequence were labeled “Disease-Relevant” if ClinVar reported a pathogenic or likely pathogenic variant at that position, and “Non-Disease-Relevant” otherwise.

### Disease-Relevancy Regressions

We utilized pre-computed residue-level evolutionary conservation scores from DescribeProt (*33* ), which calculates conservation via relative-based entropy. DescribeProt contained conservation scores for 3,015 out of 3,356 (*≈* 90%) of the Clin-Var sequences we analyzed. Residues were stratified into three disorder probability bins using IUPred2a (*9* ) scores: low ([0, 0.25)), intermediate ([0.25, 0.75)), and high ([0.75, 1.0]), and all analyses were performed separately within each bin.

To decompose the relationship between attention rollout and disease relevance, we fitted two classes of regression model within each disorder bin. First, ordinary least squares (OLS) models were fitted with attention rollout as the outcome, evaluating the attenuation of the pathogenicity coefficient (*β*_pathogenicity_) upon inclusion of conservation.

Second, logistic regression models were fitted with binary pathogenicity status as the outcome, where both predictors were *z*-scored within each bin prior to fitting.

All models used cluster-robust standard errors, clustering on protein sequence, to account for non-independence of residues within the same protein.

### Molecular Dynamics Contact Maps

Coarse-grained molecular dynamics trajectories for all intrinsically disordered regions in the human proteome were obtained from Tesei et al. (*34* ). These simulations were performed using the CALVADOS 2 coarse-grained model, which is specifically parameterized for intrinsically disordered proteins (*39* ). *Cα − Cα* distances for contact pairs were calculated for each frame using MDTraj (*40* ) with the scheme = “ca” option. A residue pair (i, j) was considered in contact if the *Cα − Cα* distance was less than 2^1*/*6^*σ_ij_*, wherein *σ_ij_* is the average of the two residues’ bead size (*41* )

The contact frequency *f_ij_*for each residue pair was calculated as: *f_ij_* = (number of frames where *d_ij_<*2^1*/*6^*σ_ij_*)*/*1000 where *d_ij_* is the *Cα − Cα* distance between residues i and j. Since all proteins were analyzed using the same number of equilibration frames (1,000), contact frequencies are directly comparable across proteins of different lengths.

These contact maps represent the fraction of time each residue pair spends in proximity within the equilibrated conformational ensemble.

### *R_g_* Multi-Layer Perceptron Training

An MLP was trained on ESM-2 final-layer mean-pooled embeddings (1280-dimensional) of sequences from the human IDRome (*34* ) to predict *R_g_*after subtracting a length-dependent baseline to encourage the model to learn sequence composition signals beyond chain length. The baseline was a second-degree polynomial fitted to log(*N* ) on training sequences only, and the model target was the residual *R_g_ − p*(log(*N* )), where *p* is the fitted polynomial. For reference, a Flory scaling fit on training data yielded *ν* = 0.533 and *R*_0_ = 2.420.

Sequences were clustered at 40% identity using CD-HIT (*24* ), and clusters were used to perform a group-stratified split into training (20,368 sequences), validation (2,075 sequences), and held-out test (5,444 sequences) sets, ensuring no sequence-similar proteins leaked across splits.

The MLP consisted of four layers of linear projection (1280 *→* 512 *→* 256 *→* 128 *→* 1), each followed by layer normalization, GELU activation, and dropout, except the final output layer. The model was trained using AdamW (learning rate 1*e −* 3, weight decay 1*e −* 4) with cosine annealing and mixed-precision training. Dropout was set to 0.3, and early stopping with patience 20 was applied based on validation loss.

### Electrostatics Zero Ablations

To assess the functional contribution of charge-sensitive attention heads to *R_g_* prediction, we performed a targeted zero-ablation analysis on ESM-2. Attention heads were ranked by their mean normalized attention to negatively charged residues (Asp, Glu) or positively charged residues (Lys, Arg), computed across a held-out set of proteins. Normalized attention was defined as the the sum of attention received at head *j* and layer *i* by each amino acid identity across residues of the sequence *p*, normalized by the occurence of the amino acid in the sequence. This value was averaged across heads, layers, and all proteins in the dataset (4,240 PDB and DisProt proteins).

Ablation was implemented by registering forward pre-hooks on the output projection of each targeted self-attention module, zeroing the pre-projection output corresponding to each selected head while leaving all other heads and model weights unchanged; this ensures that only the contribution of the targeted head, rather than its post-projection mixture with other heads, is removed. For the head-sweep analysis, at each step *n* (ranging from 1 to 50), the cumulative top-*n* ranked heads were ablated, together with an equivalent number of randomly selected heads (excluding those already targeted) ablated in parallel as a null comparator at each step. The random ablations were repeated for 5 random permutations at each *n* number of heads ablated. For all subsequent analyses, the top 50 ranked heads for each charge category were ablated jointly as a fixed set. *R_g_* predictions were generated under each ablation condition using the frozen MLP decoder, converted to absolute *R_g_* via the fitted scaling polynomial, and normalized by the Flory scaling relationship (*R_g_/*(*R*_0_*N^ν^*)); performance was evaluated against ground-truth *R_g_* values using Spearman and Pearson correlation and mean absolute error.

## Author Contributions

L.N. conceived and designed the study, developed and implemented the computational analyses, performed all analyses, and generated the figures. L.N. led the interpretation of the results and wrote the manuscript. R.S and L.L.G provided scientific guidance and contributed to interpretation of the results. T.P.J.K. supervised the study and provided guidance on study design, interpretation, and manuscript preparation. All authors reviewed and approved the final manuscript.

## Data, Code, and Materials Availability

All code, processed input datasets, figure source data, and the trained radius of gyration predictor are publicly available at https://github.com/lnaworsk/plm interp, with a stable, archived release at Zenodo (https://doi.org/10.5281/zenodo.24090068). Raw ESM-2 attention matrices, attention rollout values, and categorical Jacobian contact maps are not deposited due to size but can be regenerated using the archived code. Publicly available data were obtained from DisProt (release 2025 06, accessed 28 July 2025) (*19* ), the Protein Data Bank (accessed 16 July 2025) (*20* ), ClinVar (accessed 3 February 2026) (*32* ), NCBI Gene (*37* ) and RefSeq (GRCh38) (*38* ), IDRome (*34* ), experimental SAXS data (*35* ), and DescribeProt (*33* ).

## Supplementary Information

**Fig. S1:**
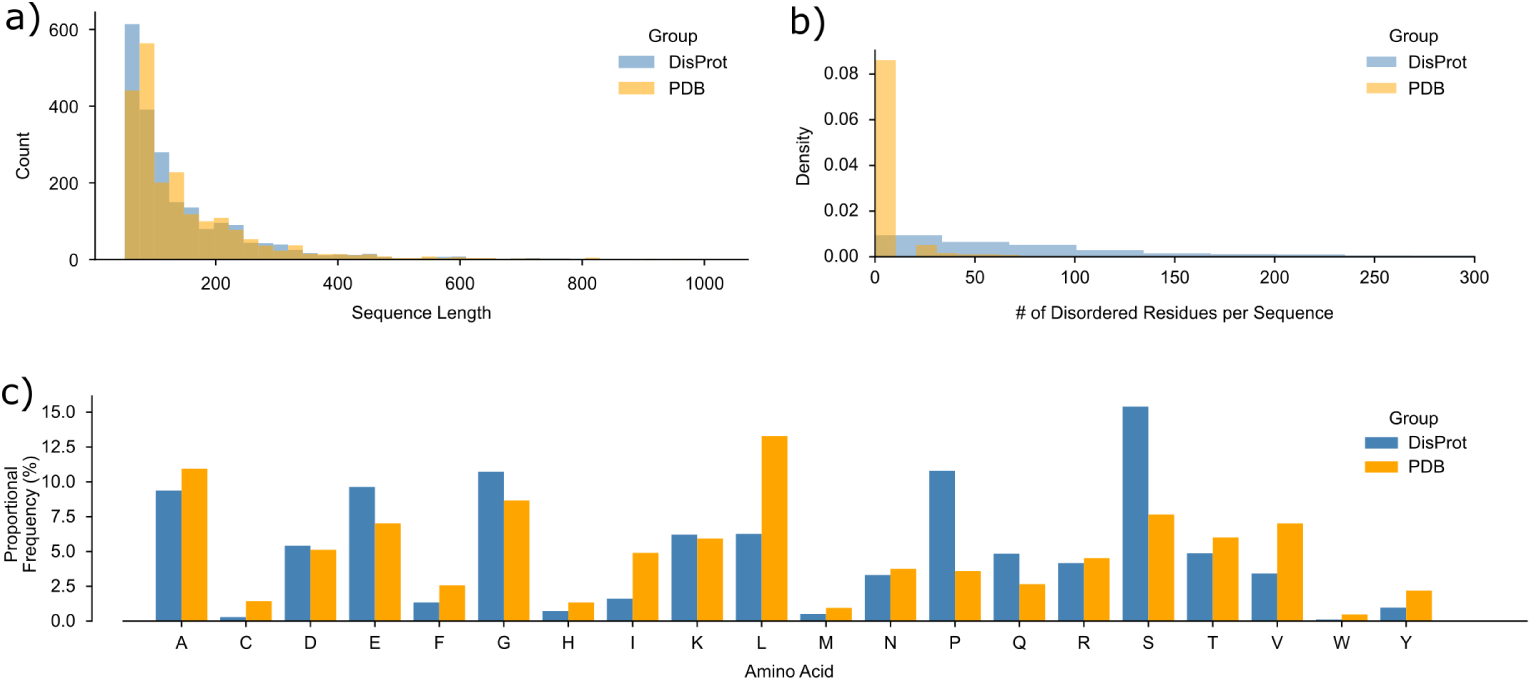
The datasets of folded proteins from PDB and IDPs from DisProt are matched for sequence length distributions, and see differences in number of disordered residues and amino acids compositions between the two groups. A) The distribution of all DisProt (blue, n = 2,120) and PDB (orange, n = 2,120) sequences’ lengths are displayed, highlighting the close similarity of sequence length distributions utilized in this study. B) The number of disordered residues per sequence (limited to up to 200 residues for visualization), as calculated by IUPred2A (*9* ), for each DisProt and PDB protein is shown, highlighting the majority of PDB sequences do not have any disordered regions, while DisProt proteins see a distribution of lengths of disordered regions. Disordered residues are counted by how many consecutive residues have a predicted disorder score above 0.5 for stretches longer than 20 residues, returning the total number of residues in those disordered regions. C) The number of occurrences of each type of amino acid across all sequences in the DisProt set, and separately, in the PDB set are divided by the total number of residues in each set, revealing proportional frequencies of each amino acid type.

**Fig. S2:**
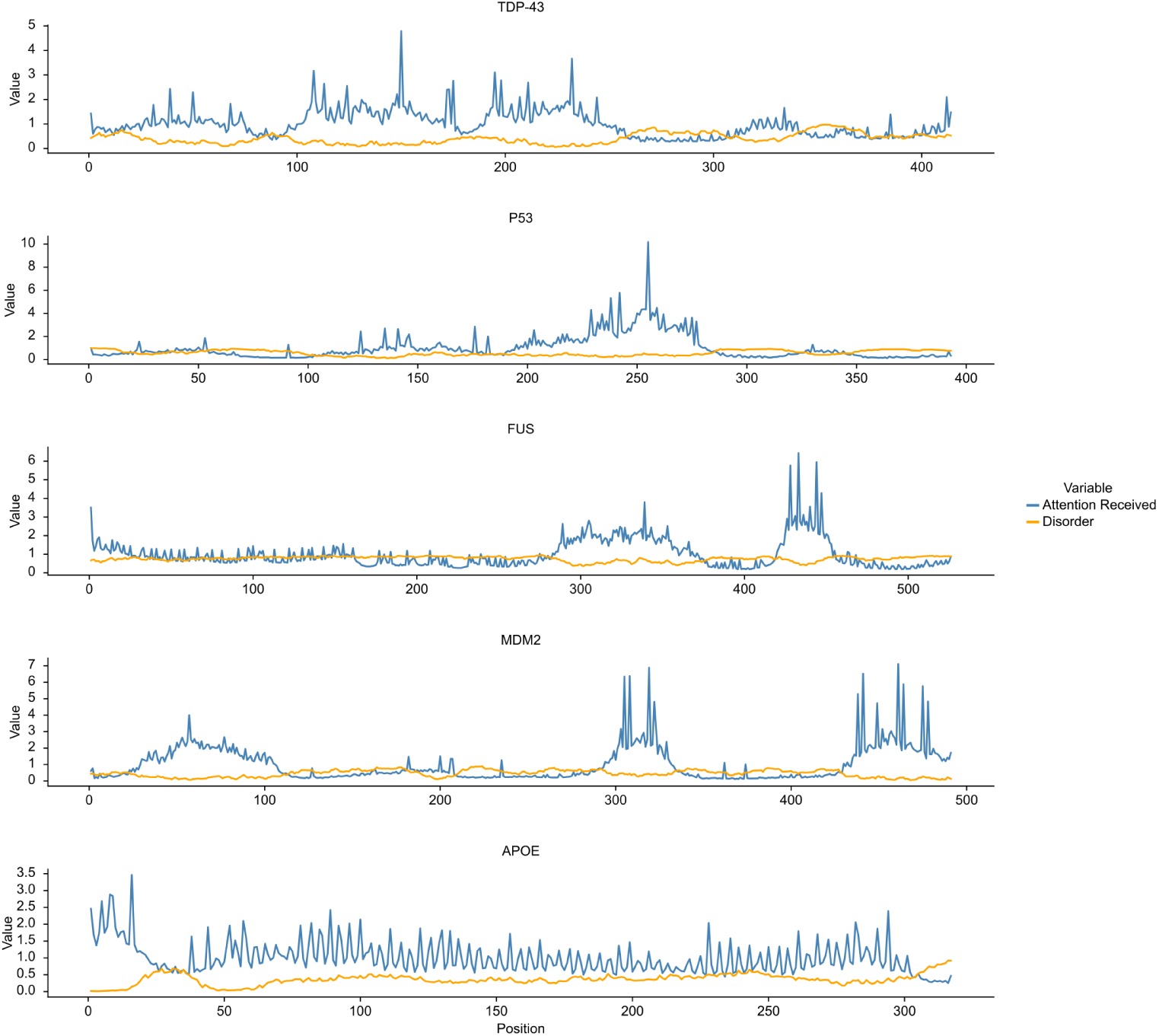
Attention rollout and disorder of example proteins. Attention rollout received and IUPred2A disorder probability across each residue in different proteins exemplifies the inverse relationship between attention received and disorder.

**Fig. S3:**
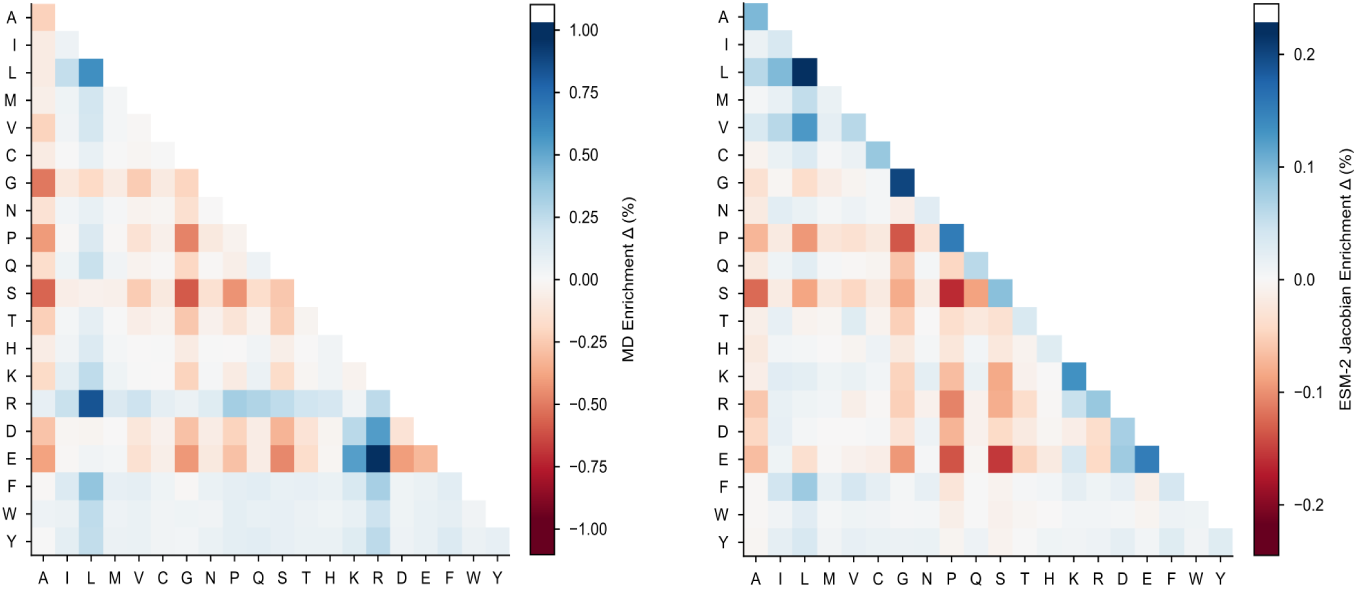
Enrichment and depletion of amino acid contact pairs in top decile relative to baseline. To compare ESM-2-derived and MD-derived contact maps, we report the baseline frequency of each amino acid pair in the top decile of each method’s (Left: MD-derived, Right: ESM-2-derived) ranked contact strenghts relative to the baseline frequency across all residues.

**Fig. S4:**
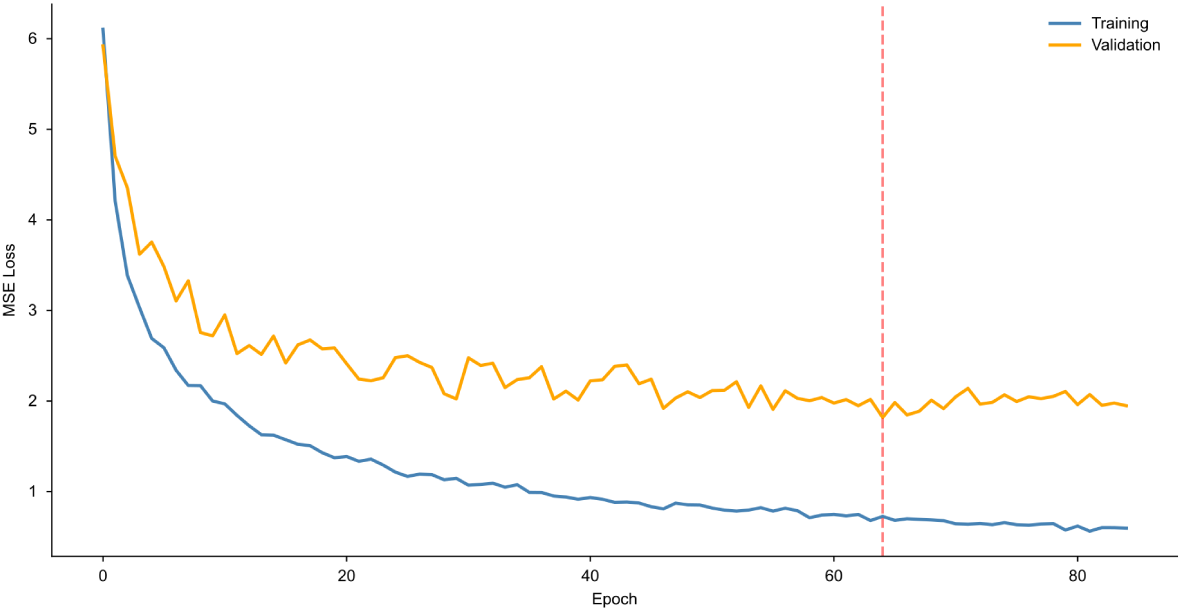
*R_g_* **MLP training and validation loss.** Training and validation loss of the MLP *R_g_* predictor is shown across each epoch, with the lowest validation loss epoch (final model) highlighted in red dashed line.

**Fig. S5:**
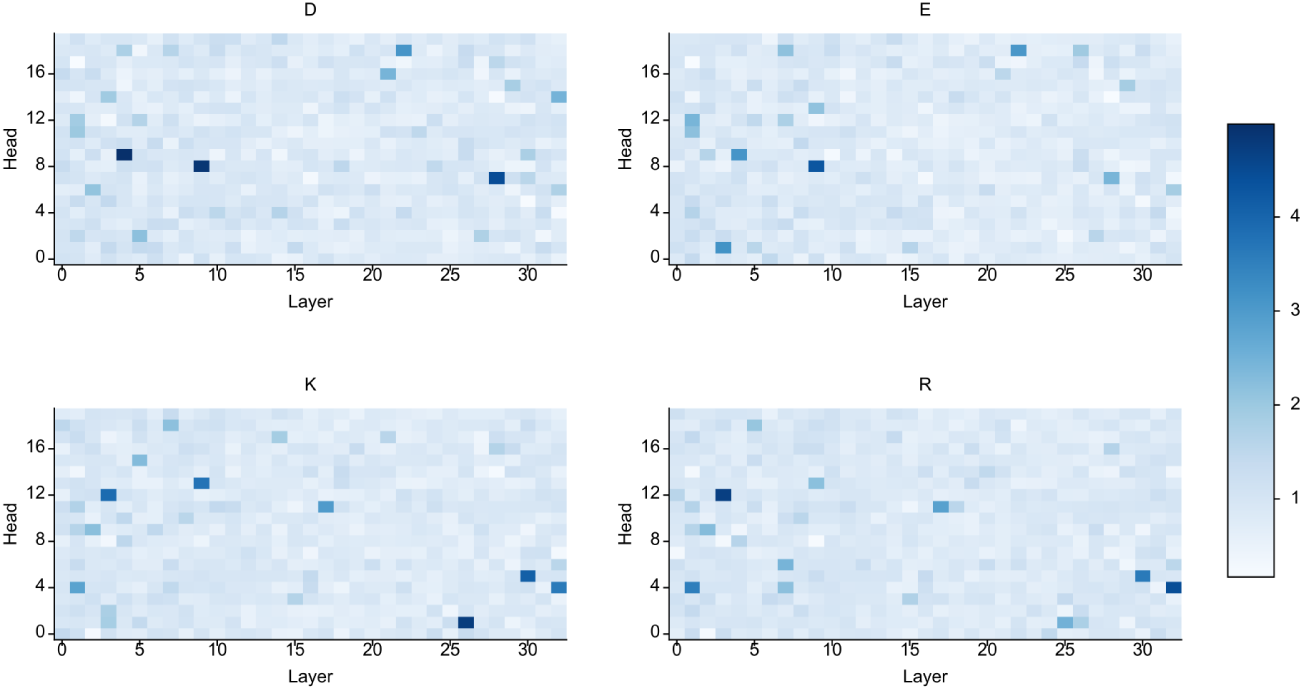
Attention allocation to charged residues. For each protein in the DisProt and PDB sets (n = 4,240), the total attention received across the sequence by Asp, Glu, Lys, and Arg residues separately was summed and normalized by occurrence in the protein. This value was then averaged across all proteins at each head and layer.

